# Degeneracy in the neurological model of auditory speech repetition

**DOI:** 10.1101/2022.03.25.485823

**Authors:** Noor Sajid, Andrea Gajardo-Vidal, Justyna O. Ekert, Diego L. Lorca-Puls, PLORAS team, Thomas M. H. Hope, David W. Green, Karl J. Friston, Cathy J. Price

## Abstract

In the neurological model of language, repeating heard speech involves four left hemisphere regions: primary auditory cortex for processing sounds; Wernicke’s area for processing auditory images of speech; Broca’s area for processing motor images of speech; and primary motor cortex for overt speech articulation. Previous functional-MRI (fMRI) studies confirm that auditory repetition activates these regions. Here, we used dynamic causal modelling (DCM) to test how the four regions interact with each other during single word and pseudoword auditory repetition. Contrary to expectation, we found that, for both word and pseudoword repetition, the effective connectivity between Wernicke’s and Broca’s areas was predominantly bidirectional and inhibitory; activity in the motor cortex could be driven by either Wernicke’s area or Broca’s area; and the latter effect varied both within and between individuals. Such variability speaks to degenerate functional architectures that support auditory repetition and may explain resilience to functional loss after brain damage.

## Introduction

Auditory speech repetition involves the immediate reproduction of heard speech. It requires the successful translation of auditory input into a motor output that matches the heard speech. The influential 19th century neurological model of language ^1, 2, 3, 4^, later refined by Norman Geschwind ^4^, posited that speech repetition involves a sequential flow of information across four left hemisphere brain regions: the primary auditory cortex, the left posterior superior temporal cortex (Wernicke’s area), the left posterior inferior frontal gyrus (Broca’s area) and the primary motor cortex, with information being relayed from Wernicke’s to Broca’s areas via the arcuate fasciculus. More recent studies have challenged this model of the functional anatomy of language. For example, the brain regions activated during speech repetition include multiple cortical and subcortical areas that are not part of the neurological model ^5^. Likewise, white matter tracts, other than the arcuate fasciculus, have been shown to connect temporal and frontal regions ^6, 7^. These and other observations led Tremblay and Dick ^8^ to claim that the “Wernicke-Lichtheim-Geschwind” model is obsolete, in line with contemporary demonstrations that other (indirect) pathways might support auditory word repetition ^9, 10^.

Despite widespread documentation that the neurological model of language is over-simplified, there is strong neuroimaging evidence^5^ that auditory speech repetition activates regions that fall in the vicinity of Wernicke’s and Broca’s areas. Specifically, the posterior superior temporal sulcus (pSTS) and pars opercularis (pOp) were selected as proxies; guided by fMRI findings from an independent group of 25 neurologically intact participants who performed the same word and pseudoword repetition tasks ^5^. Thus, one can argue that the neurological model captures the core of the auditory speech repetition system when its components are precisely localised within pSTS and pOp, while acknowledging that repetition also involves other cortical and subcortical regions.

Here, we asked how activity within the primary auditory cortex (A1), pSTS, pOp or primary motor cortex (M1) influences each of the remaining three areas during auditory word and pseudoword repetition using Dynamic Causal Modelling (DCM) ^11, 12, 13, 14^, of functional MRI data from 59 healthy subjects. Using DCM, we estimated the effective connectivity across the four left hemisphere regions (A1, pSTS, pOp and M1) during auditory speech repetition, and evaluated the evidence for the neurological model.

If the neurological model is correct, the results should show that activity in the primary auditory cortex excites pSTS, pSTS excites pOp and pOp excites the primary motor cortex. Evidence for this model was compared against evidence for alternative models that allowed primary motor cortex to be driven by Wernicke’s area as well as Broca’s area or instead of Broca’s area and (b) inhibitory as well as excitatory extrinsic (i.e., between-region) connectivity. By limiting our study to the four regions comprising the neurological model, we were able to address our hypotheses simply and directly, while remaining agnostic as to the exact white matter tracts that support the directed interactions (i.e., effective connectivity) between brain regions. For example, effective connectivity between pOp and motor cortex does not imply monosynaptic connections via a direct white matter tract; rather, polysynaptic connections involving one or more fasciculi. In other words, effective connectivity can be mediated vicariously through intervening cortical stations (not included in the model).

We were particularly interested in the evidence for degenerate functional architectures; namely, whether effective connectivity varies across participants (inter-subject variability) or within participants (intra-subject variability) for different tasks (word or pseudoword). To ensure that inter-subject variability was not a consequence of the precise anatomical definition of regions, we compared effective connectivity using two subregions of pOp (i.e., dorsal and ventral) and two subregions for motor control (i.e., face and tongue with larynx). Characterising potentially degenerate architectures, in terms of variation within and between subjects, might provide insights into how language functions recover following neurological damage. Under this formulation, structures (e.g., subgraphs of a neuronal network) may be sufficient, but not necessary, for a particular function—meaning that functional deficits arise only when all degenerate structures are damaged ^15, 16, 17^.

## Results

### Group-level effective connectivity for word and pseudoword repetition

The group-level effective connectivity estimates for word repetition are presented in Figure 1 and Table 1. Extrinsic (between-region) connections are parameterised directly as rate constants (i.e., the rate of change in a target region, per unit change in the source region). In contrast, intrinsic (self) connections in DCM for fMRI are log-scaling parameters that are applied to inhibitory connections to ensure dynamical stability. This means that a positive self-connection means greater self-inhibition

**Table 1.** Estimated connections at the group level for word and pseudoword repetition for the subregional configurations considered. Extrinsic connections are in units of hertz (Hz) because they are rates of change (rate constants). Only estimated connections with a posterior probability greater > 0.75 are shown. Here, 1-4 denotes the different subregional configurations: 1 (M1-f and dpOp), 2 (M1-f and vpOp), 3 (M1-tl and dpOp) and 4 (M1-tl and dpOp).

| *From* | *To* | Word repetition |  |  |  | Pseudoword repetition |  |  |  |
| --- | --- | --- | --- | --- | --- | --- | --- | --- | --- |
|  |  | 1 | 2 | 3 | 4 | 1 | 2 | 3 | 4 |
| *A1* | A1 | - | - | -0.09 | - | 0.07 | 0.15 | - | - |
| *A1* | pSTS | 0.51 | 0.49 | 0.42 | 0.42 | 0.53 | 0.52 | 0.33 | 0.35 |
| *A1* | pOp | 0.42 | 0.39 | 0.31 | 0.32 | 0.43 | 0.24 | 0.33 | 0.20 |
| *pSTS* | A1 | -0.28 | -0.36 | -0.44 | -0.35 | - | -0.09 | -0.15 | - |
| *pSTS* | pSTS | - | -0.15 | -0.18 | -0.10 | -0.18 | -0.10 | -0.37 | -0.43 |
| *pSTS* | pOp | -0.27 | -0.18 | -0.25 | -0.20 | - | -0.20 | -0.14 | -0.13 |
| *pSTS* | M1 | 0.66 | 0.88 | 0.38 | 0.41 | 0.53 | 0.73 | 0.27 | 0.38 |
| *pOp* | A1 | - | - | - | - | - | -0.19 | -0.25 | -0.26 |
| *pOp* | pSTS | -0.22 | -0.21 | -0.16 | -0.18 | -0.17 | -0.27 | -0.08 | -0.18 |
| *pOp* | pOp | -0.49 | -0.40 | -0.76 | -0.63 | -0.32 | -0.58 | -0.48 | -0.54 |
| *pOp* | M1 | 0.31 | 0.13 | 0.18 | 0.19 | 0.32 | - | 0.15 | 0.06 |
| *M1* | A1 | -0.40 | -0.29 | -0.38 | -0.33 | -0.30 | -0.20 | -0.14 | -0.25 |
| *M1* | pSTS | -0.11 | -0.12 | -0.13 | -0.04 | -0.19 | -0.10 | -0.16 | -0.09 |
| *M1* | pOp | -0.13 | -0.10 | -0.13 | -0.10 | -0.21 | - | -0.15 | 0.05 |
| *M1* | M1 | 0.41 | 0.52 | 0.09 | 0.15 | 0.31 | 0.20 | -0.26 | -0.13 |

**Figure 1.**
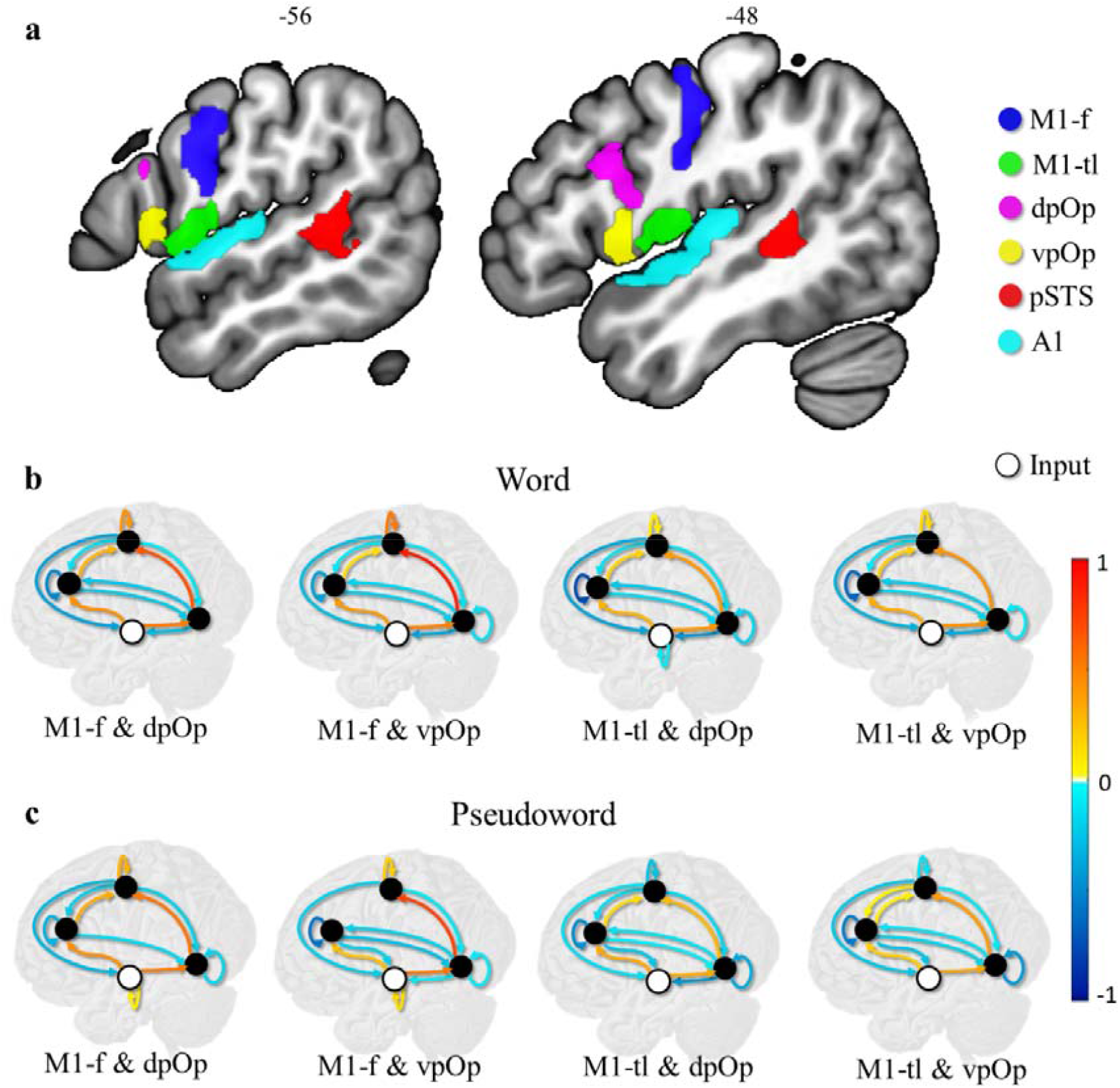
Group level DCM model after Bayesian model selection. The top row shows the 6 anatomical regions of interest: M1-f (blue), M1-tl (green), dpOp ^18^, vpOp (yellow), pSTS (red) and A1 (cyan). The middle row shows the strength of effective connectivity among regions, during word repetition. In each model, the four circles represent A1 (bottom, white circle, input area), pSTS (black circle, right), pOp (black circle, left) and M1 (black circle, top). Four different models are depicted for either vpOp or dpOp; with either M1-f or M-tl. The third row shows the same for pseudoword repetition. Red lines denote positive connections of 1, dark blue denotes negative connections of -1 and the rest represent gradation between the two. Positive extrinsic connections between regions are excitatory while intrinsic self-connections are log scale parameters (that scale an inhibitory self-connection). Only estimated connections with a high posterior probability (i.e., > 0.75 representative of strong Bayesian evidence) are shown.

We observed excitatory extrinsic connectivity (i.e., activity in one region increases activity in another) from A1 to pSTS, pSTS to M1, A1 to pOp, and pOp to M1. In addition, there were inhibitory effective connections (i.e., activity in one region suppresses activity in another) from M1 to A1, pSTS to A1, and most surprisingly, between pSTS and pOp in both directions. The same results were observed for different subregional configurations: i.e., replacing the dorsal pOp (dpOp) subregion with the ventral pOp (vpOp) subregion and/or the face motor control (M1-f) subregion with the tongue and larynx motor control (M1-tl) subregion (Figure 1b). The main difference between subregions was an increased self-inhibition for pSTS for M1-f compared to M1-tf.

The estimated group-level connectivity for pseudoword repetition (Figure 1c) was very similar to the effective connectivity observed for word repetition. For example, there was an inhibitory connection from pOp (both dorsal and ventral) to pSTS during both auditory pseudoword and word repetition. The differences in effectivity connectivity for pseudoword repetition, compared to word repetition, were: (i) a negative (i.e., less inhibitory) self-connection for pSTS (ii) a negative (i.e., less inhibitory) self-connection for M1-tl, and (iii) no inhibitory connection from pSTS to A1 and from pSTS to dpOp for particular subregional configurations.

### Variation in connectivity from pOp and pSTS to M1

The neurological model suggests excitatory connections from pSTS to pOp, and from pOp to M1 but not directly from pSTS to M1. To investigate the (unanticipated) group-level effective connectivity from pSTS to M1 in addition to pOp to M1 (Figure 1), we evaluated participant specific effectivity connectivity from pOp and pSTS to M1 (Figure 2; Supplementary Figure 1).

**Figure 2.**
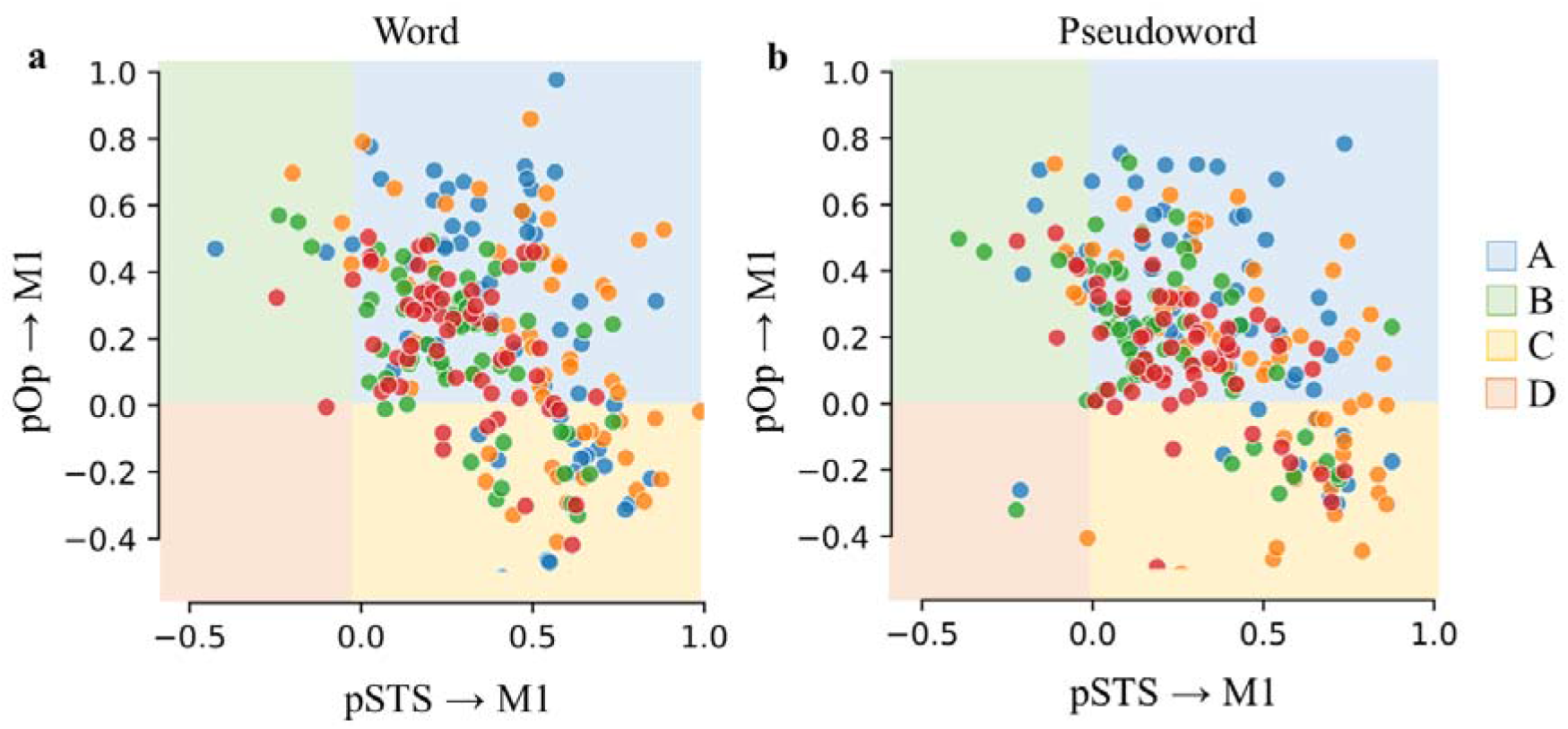
Individual-level effective connectivity from pOp and pSTS to M1. The two panels report the group membership for word (a) and pseudoword (b) repetition. Here, Group A was defined by excitatory connections from both pOp and pSTS to M1; Group B was defined by excitatory connections from pSTS to M1 but not pOp to M1; Group C was defined by excitatory connections from pOp to M1 but not from pSTS to M1; and Group D was defined by the absence of connections from both pOp and pSTS to M1 (please see Figure for colour codes).

We considered 8 combinations of subregions and task for each participant i.e., 2 M1 regions (M1-f or M1-tl) x 2 pOp regions (dpOp or vpOp) x 2 repetition tasks (Table 2). Here, each model was assigned to one of four groups (Figure 2): Group A was defined by excitatory connections from both pOp and pSTS to M1; Group B was defined by excitatory connections from pSTS to M1 but not from pOp to M1; Group C was defined by excitatory connections from pOp to M1 but not from pSTS to M1; and Group D was defined by the absence of definitive connections from both pOp and pSTS to M1. The presence of a connection was assessed by Bayesian model comparison – briefly, we compared the model evidence with and without each connection.

**Table 2.**
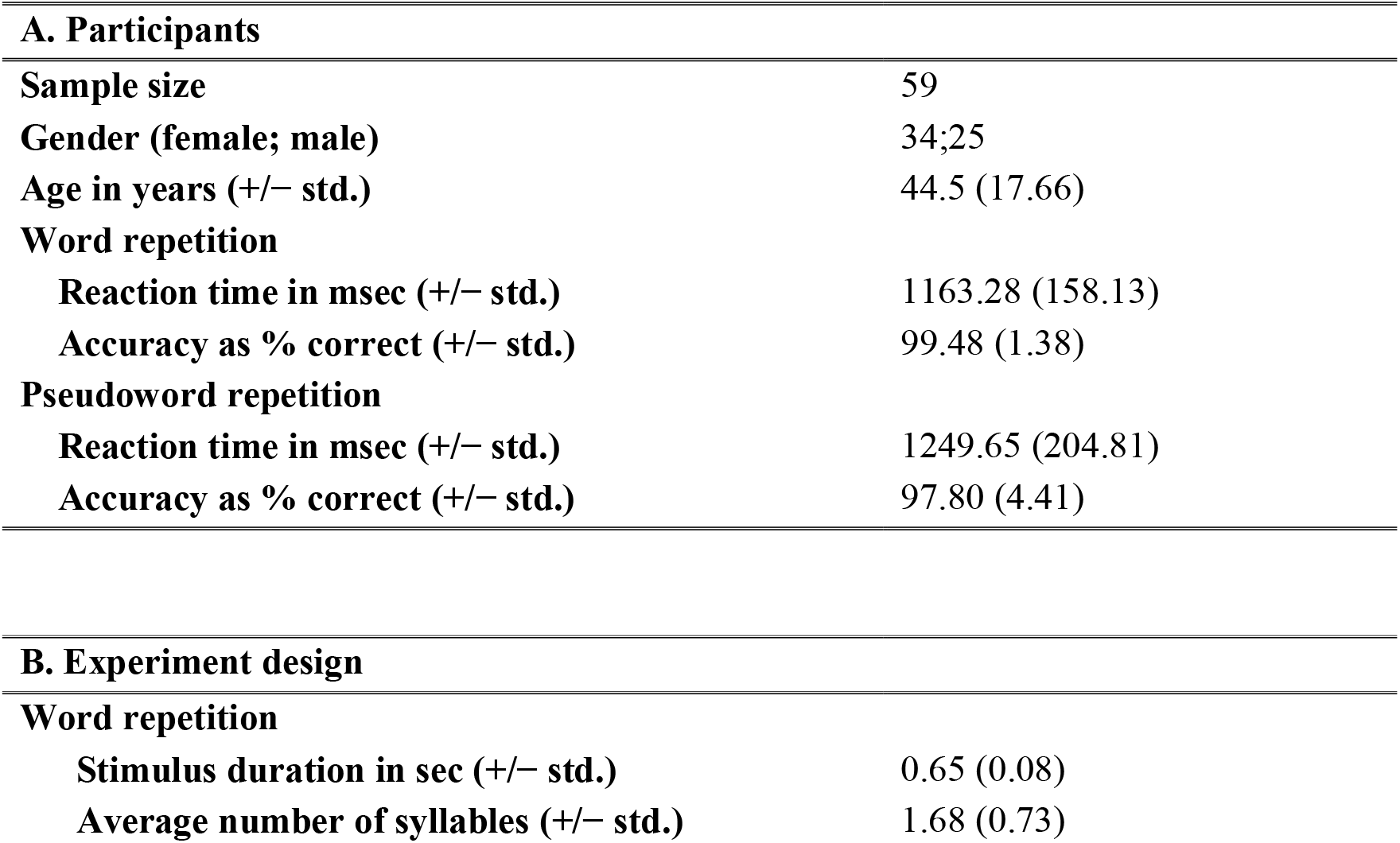
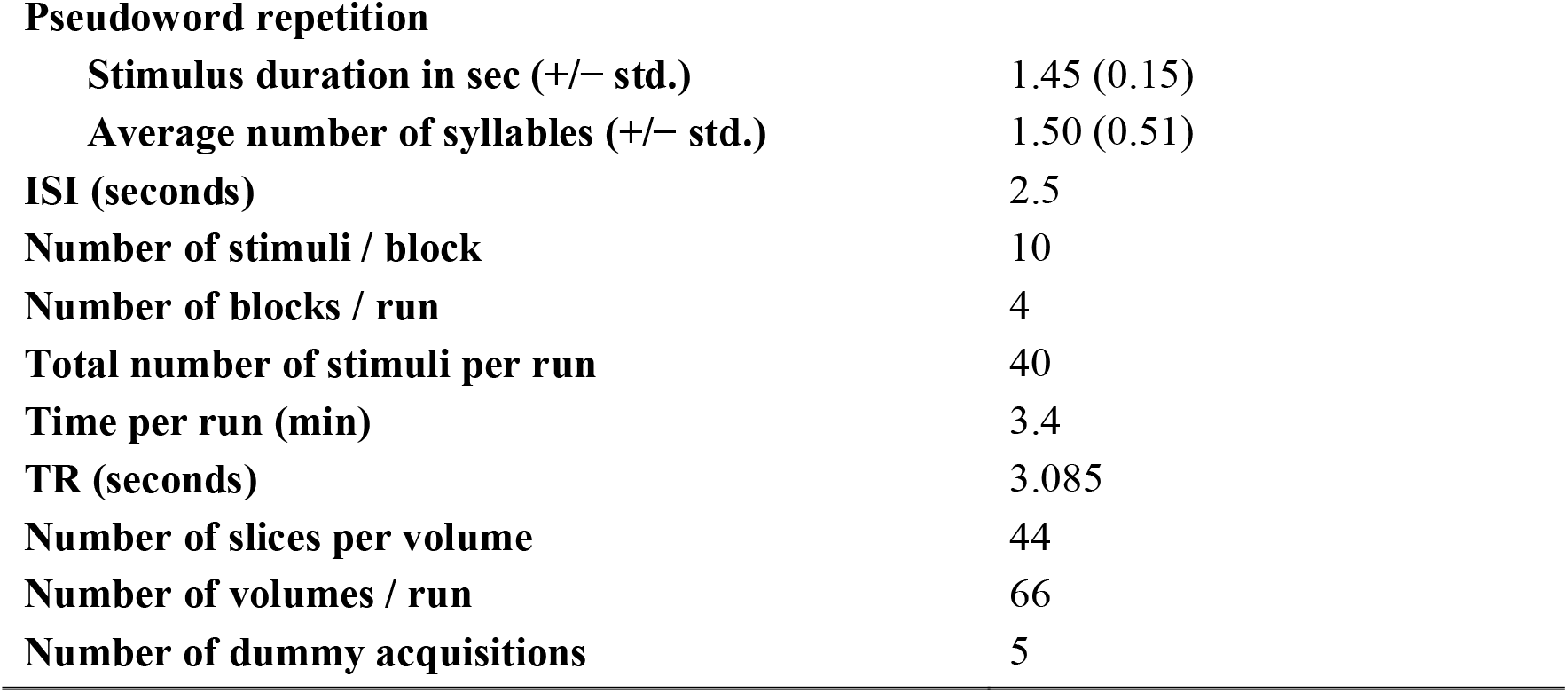
Summary of participant data (A) and experimental design specifications (B)

| **A. Participants** |  |
| --- | --- |
| **Sample size** | 59 |
| **Gender (female; male)** | 34;25 |
| **Age in years (+/- std.)** | 44.5 (17.66) |
| **Word repetition** |  |
| **Reaction time in msec (+/- std.)** | 1163.28 (158.13) |
| **Accuracy as % correct (+/- std.)** | 99.48 (1.38) |
| **Pseudoword repetition** |  |
| **Reaction time in msec (+/- std.)** | 1249.65 (204.81) |
| **Accuracy as % correct (+/- std.)** | 97.80 (4.41) |
| **B. Experiment design** |  |
| **Word repetition** |  |
| **Stimulus duration in sec (+/- std.)** | 0.65 (0.08) |
| **Average number of syllables (+/- std.)** | 1.68 (0.73) |
| **Pseudoword repetition** |  |
| **Stimulus duration in sec (+/- std.)** | 1.45 (0.15) |
| **Average number of syllables (+/- std.)** | 1.50 (0.51) |
| **ISI (seconds)** | 2.5 |
| **Number of stimuli / block** | 10 |
| **Number of blocks / run** | 4 |
| **Total number of stimuli per run** | 40 |
| **Time per run (min)** | 3.4 |
| **TR (seconds)** | 3.085 |
| **Number of slices per volume** | 44 |
| **Number of volumes / run** | 66 |
| **Number of dummy acquisitions** | 5 |

Across participants, more than half the models had excitatory connections from both pOp and pSTS to M1 (i.e., Group A; Figure 3a). Those with excitatory connections from pSTS to M1 but not from pOp to M1 (Group B) and those without connections from either pOp or pSTS to M1 (Group D) accounted for a further ~20% of estimated models. Importantly, <5% of estimated models fell in Group C (i.e., excitatory connections from pOp to M1 but not from pSTS to M1). Moreover, only 2% of models (10 in total) were consistent with the neurological model, i.e., Group C (pSTS->pOp->M1) with excitatory connections from pSTS to pOp, see Figure 3b.

**Figure 3.**
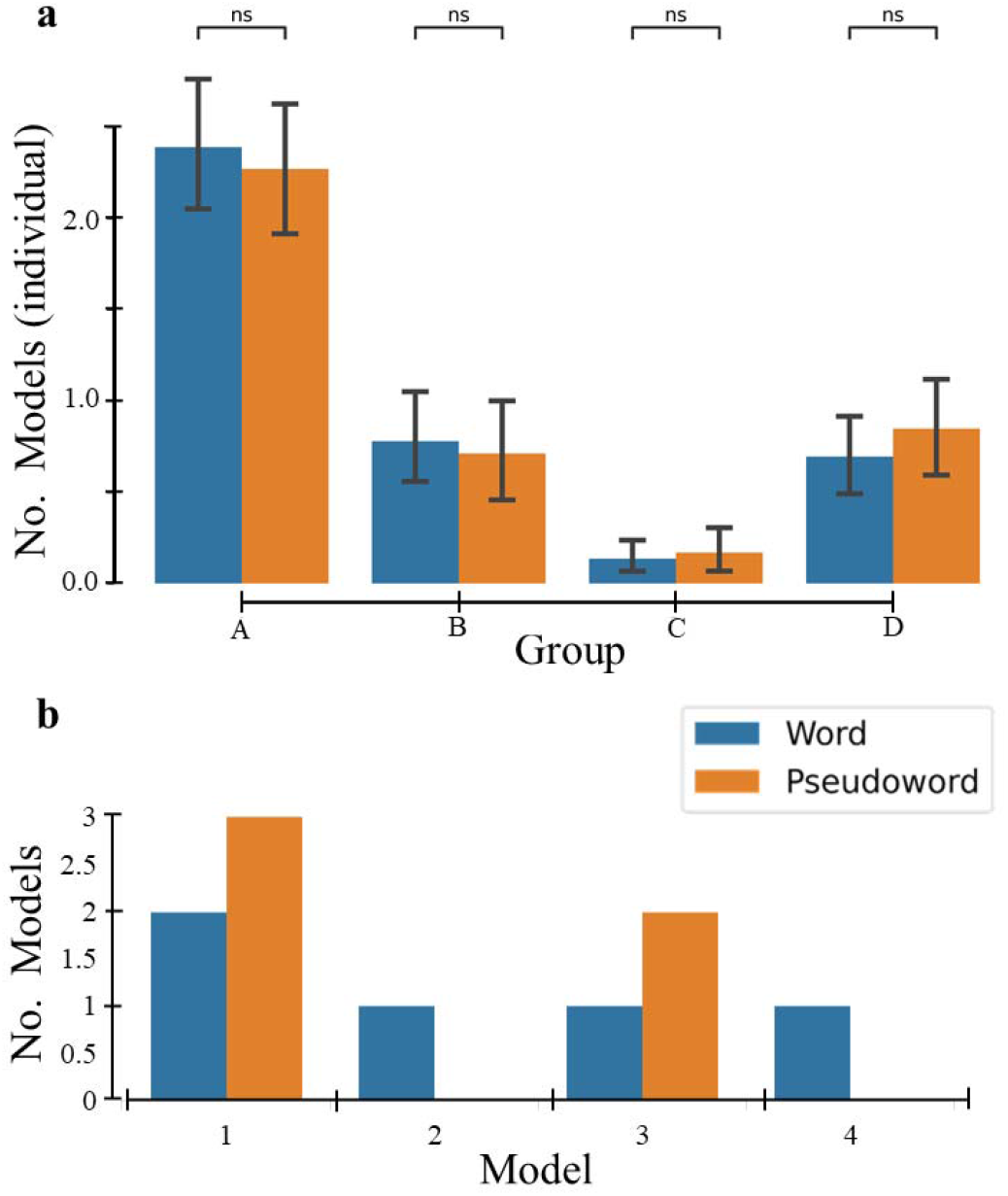
Variable group membership. (a) plots the number of models (y axis) assigned to each group (x axis), averaged across the 8 configurations i.e., 2 M1 (M1-f or M1-tl) regions x 2 pOp regions (dpOp or vpOp) x 2 repetition tasks. A denotes excitatory connections from pOp to M1 and pSTS to M1; B denotes excitatory connections from pSTS to M1 but not from pOp to M1; C denotes excitatory connections from pOp to M1 but not from pSTS to M1; and D denotes no connectivity from both pOp and pSTS to M1. We found no significant [ns] differences (with a p-value of 1.00) between word and pseudoword repetition across the different groups using a two-sided Mann-Whitney-Wilcoxon test with Bonferroni correction. (b) represents the 10 models (subset of Group C) that were consistent with the neurological model for each subregional configuration (i.e., estimated model): 1 (M1-f and dpOp), 2 (M1-f and vpOp), 3 (M1-tl and dpOp) and 4 (M1-tl and dpOp).

Concerning which M1 or pOp subregion was used, we found no significant differences in the proportion of A, B, C or D models for (i) word versus pseudoword repetition (Figure 3a) and (ii) subregional configurations including activity from vpOp versus dpOp or from face (M1-f) versus the larynx and tongue (M1-tl) (Supplementary Figure 2) using Mann-Whitney-Wilcoxon test two-sided with Bonferroni correction. In addition, we did not detect group differences in the degree of activation in any of the subregions (Supplementary Figure 3) and no group differences in behavioural performance across different subregional configuration (Supplementary Table 2).

### Degeneracy in auditory repetition

Degeneracy pertains to how many structures can be recruited to subserve a function, for example, to repeat heard speech. In our results, degeneracy is implied by the high inter- and intra-subject variability observed in group membership, i.e., variability in the functional architectures engaged by word repetition. Specifically, participants showed excitatory connectivity to M1 from pSTS, pOp or both pSTS and pOp. We assessed the variability in these connectivity architectures using the entropy of their sample density (i.e., number of observed groups per participant)^19^. Variations in effective connectivity (i.e., degeneracy) are captured by this measure ^20^.

We found that the average entropy in individual group membership for word repetition was 0.49, and in pseudoword repetition was 0.55 (Figure 4). This speaks to degeneracy in the functional architectures – and provides a novel perspective on the neurological model of auditory speech repetition. Our measure of degeneracy indicates that individuals, in our sample, are likely to be able to execute the repetition tasks in different ways: e.g., for word repetition, subject C073 belonged to both Group B (3x) and Group C (1x). See Supplementary Table 1 for a breakdown of how groupings differed across the different subregional configurations. Importantly, this intra-participant variability, and the similarity of the models we observe at the group level for words and pseudowords, makes it highly unlikely that inter-participant variability can be explained solely in terms of between-participant variability in brain structure and functional anatomy.

**Figure 4.**
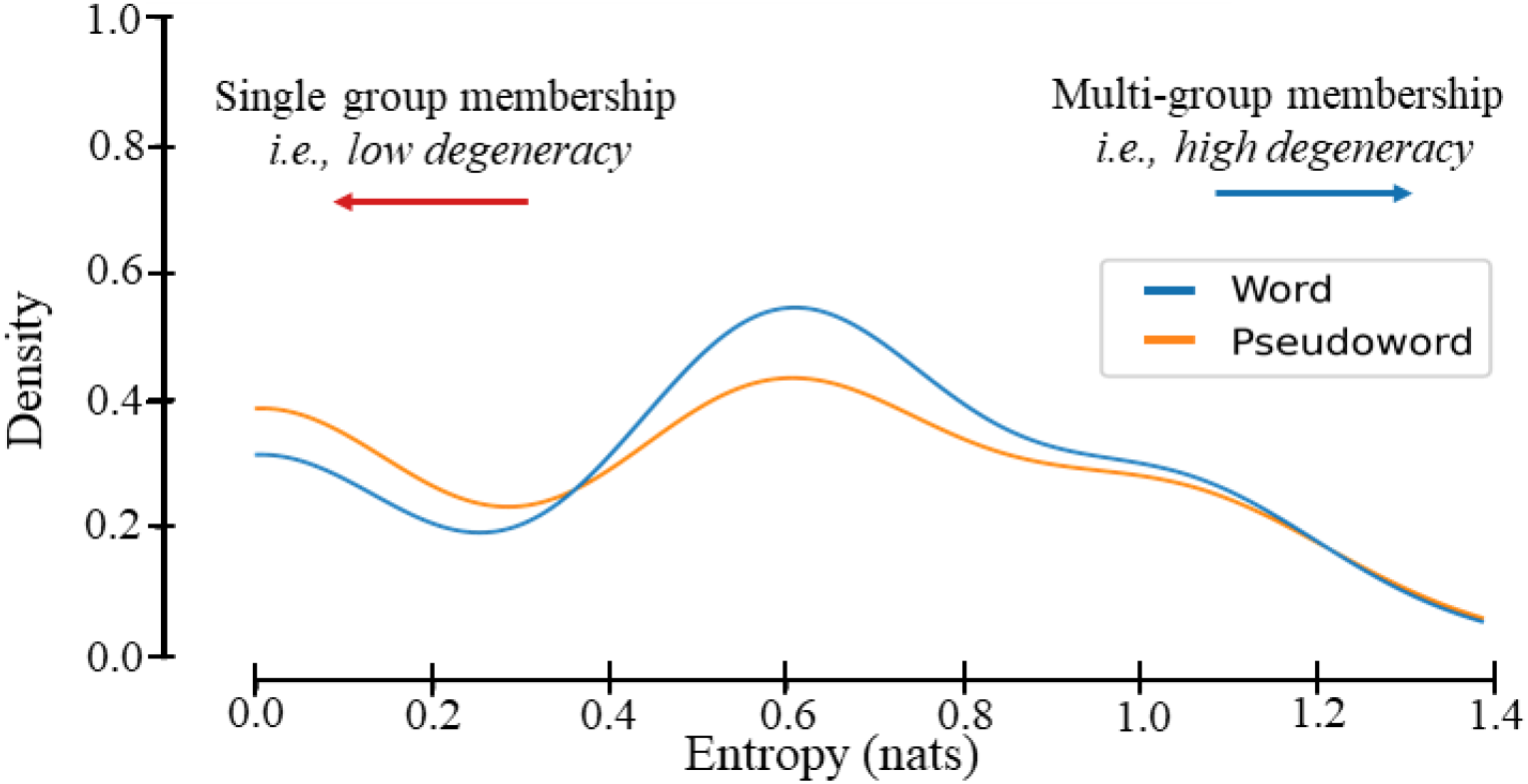
Degenerate group membership. Plot of the sample density estimate (y axis) of the dispersion (i.e., entropy, x axis) over group membership for word (blue) and pseudoword (orange) repetition. Here low entropy (=0; red arrow) denotes membership of a single group (e.g., the participant was consistently in group A) whereas high entropy (>1.3; blue arrow) denotes membership dispersed over all groups (i.e., the participant was in group A, B, C and D for the different subregional combinations).

## Discussion

The neurological model of language ^1, 2, 3, 4^ posits that, during speech repetition, information flows sequentially from A1 to Wernicke’s area, Wernicke’s area to Broca’s area (via the arcuate fasciculus), and finally Broca’s area to M1. In this study, we evaluated the evidence for this hypothesis by modelling the effective connectivity among these regions using DCM of fMRI data from healthy participants. Briefly, our results show, in contradiction with the neurological model, that pSTS (as a proxy for Wernicke’s area) drives M1 independently of pOp (as a proxy for Broca’s area). Together with the existence of inhibitory connectivity between pOp and pSTS, and evidence of both inter-participant and intra-participant variability, our study demonstrates a distributed, functional heterarchy along the (sub) regions of interest: A1, pSTS, pOp and M1 during word and pseudoword repetition. Such variability is indicative of degeneracy in the ways words and pseudowords can be repeated and has potentially important implications for functional recovery after brain damage. This motivates further experiments to find the grey matter regions and white matter tracts that underlie the effective connectivity from Wernicke’s area to the primary motor cortex, bypassing Broca’s area.

At a general level, our DCM results show that pSTS and pOp sit at the same level of the cortical hierarchy for speech repetition, with A1 below and M1 above. We briefly justify this interpretation. It is based on the pattern of excitatory and inhibitory connections, under the predictive processing assumption that forward connections (up the hierarchy) have to be excitatory, and backward connections (down the hierarchy) are inhibitory or less excitatory ^21, 22, 23^. More specifically, excitatory effective connectivity reflects prediction errors being passed from a lower to a higher area whereas inhibitory effective connectivity reflects the explaining away (reduced excitation) of the prediction error lower in the hierarchy, generally thought to be mediated by inhibitory interneurons within the target region ^24, 25, 26, 27^.

The neurological model predicts that during word and/or pseudoword repetition, pSTS (as a proxy for Wernicke’s area) would interact directly with pOp (as a proxy for Broca’s area), and indirectly with M1 through pOp. In agreement with this model, we found that, during word repetition, there were excitatory forward connections from pOp to M1 at the group level. However, at the individual level this effective connection was inconsistent. Contrary to the predictions of the neurological model, we found that during word/pseudoword repetition (i) pSTS exerts an excitatory influence on M1 that cannot be explained by afferents from pOp, and this effective connection was more consistently observed across participants than the excitatory effective connectivity from pOp to M1; (ii) the connections between pSTS and pOp were inhibitory rather than excitatory; (iii) there were profound inter-participant variations in whether M1 was driven by pOp, pSTS or both; and (iv) the intra-subject variability in effective M1 afferents depended on whether: the stimuli were words or pseudowords, the ventral or dorsal aspect of pOp region was considered; or the M1 region corresponded to the face or tongue and larynx area.

Despite the individual variability, the group-level results were remarkably similar for pseudoword repetition even when activity was taken from different parts of M1 or pOp. One exception was that, unlike during word repetition, we did not observe inhibitory connections from pSTS to A1 for particular subregional configurations. Speculatively, under a predictive coding account, there may be lower precision in predictions for auditory processing when the stimuli are always unfamiliar within a run (i.e., the pseudoword condition) compared to when the stimuli are always familiar within a run (i.e., the word condition).

Below, we discuss the implications of each finding (connectivity from pSTS to M1; bidirectional inhibition between pSTS and pOp and inter- and intra-participant variability in effective connectivity) along with necessary directions for future experiments.

### Connectivity from pSTS

Positive (excitatory) connections from pSTS to M1 were observed during word repetition (in all but 3 participants) and in pseudoword repetition (in all but 8 participants). This effective connection is not consistent with the predictions of the neurological model and cannot be explained by lack of activity in pOp, which was consistently activated across participants—with most showing excitatory connectivity from pOp to M1: 39/59 during word repetition and 44/59 during pseudoword repetition. It was also surprising that connectivity from pSTS to pOp was inhibitory. Together, these observations provide evidence against the assumption of the neurological model that pOp mediates the influence of pSTS on M1.

Although not consistent with the neurological model, the separable influence of pSTS and pOp on M1 might explain (at least partially) why focal damage to pOp does not result in long-lasting speech production impairments ^28, 29, 30^. Specifically, the connectivity from pSTS to M1 might be one of the key mechanisms underlying speech production recovery after pOp damage. Future fMRI studies of patients with relatively focal pOp damage^31^ could therefore investigate the degree to which the effective connectivity from pSTS to M1 is related to preserved speech production abilities.

Our study does not elucidate which anatomical pathways sustain neuronal message passing from pSTS to M1. Theoretically, the supporting pathways may lie dorsal or ventral to the Sylvian fissure ^32^ and different pathways may be required for word and pseudoword repetition ^33^. Previous studies have demonstrated a division of labour between the ventral and dorsal processing routes underlying auditory word repetition ^32, 33, 34, 35^. A dorsal parietal-frontal stream is proposed to support form-to-articulation mapping and a ventral temporal– frontal stream is proposed to support form-to-.meaning mapping ^36^. However, this division of labour is not sufficient to explain our findings during pseudoword repetition that cannot be supported by mapping form-to-meaning. Future investigations are therefore required to establish how different pathways drive motor activity in M1.

### Bidirectional inhibition between pSTS and pOp

The bidirectional inhibition between pOp and pSTS in the cortical hierarchy may mediate a turn-taking relation between them. Given that pSTS is strongly associated with speech perception and pOp is strongly associated with the encoding of a speech plan (in both the 19th-century neurological model and 21st-century neuroscience), turn-taking between pSTS and pOp is consistent with perceiving a word before repeating it. In other words, we expect reciprocal inhibition in terms of inferring what has been heard (excitation in pSTS) and what the participant is saying (excitation in pOp). This mutual inhibition is further endorsed by the phenomena of sensory attenuation; namely, the attenuation of self-produced sensations during speech ^37, 38, 39^.

### Inter-participant and intra-participant variability in the connections to M1

Inter-participant variability analysis considered three distinct hierarchical structures. The most common functional architecture (Group A) was a heterarchical organisation where pSTS and pOp can be thought of as superordinate hierarchically to A1, with no clear hierarchical relationship between themselves. Conversely, Group B and Group C conformed to a hierarchical organisation. For Group B, this took the form of excitatory efferent connections from A1 to pSTS, and from pSTS to M1 and pSTS to pOp. Additionally, Group B featured inhibitory afferent connections from pOp to pSTS. From this, we infer that, under a predictive processing account, pSTS is lower in the functional hierarchy than pOp for Group B. In contrast, for Group C, we observe the opposite pattern where excitatory efferent connections are from A1 to pOp, pOp to M1, and pOp to pSTS: i.e., pSTS is hierarchically superordinate to pOp. For Group C, a very small subset (2%) featured connectivity consistent with the classic pathway: i.e., excitatory connectivity from pSTS to pOp, and pOp to M1 but not pSTS to M1.

However, even here, the use of the classic pathway varied across task and subregional configurations. These distinct hierarchies are examples of degenerate functional architectures underwriting word repetition ^16, 20, 40^. Participants might repeat words/pseudowords by either engaging pOp or using an alternative pathway involving pSTS. Moreover, the fact that we observed the same participants moving from one hierarchical functional architecture to another, when repeating words or pseudowords, provides evidence that they were able to engage more than one processing route, whereas others prefer one over another, contingent on the experimental condition (word or pseudoword repetition).

This intra-participant variability has implications for how lesions to pSTS or pOp might change the effective connectivity of the network. Contrary to the neurological model, but consistent with current studies ^28^, we expect disconnections between pOp and pSTS—as a result of direct damage to pOp—to induce transitory auditory word repetition deficits. This is because damaging either region would mediate a readjustment of the overall effective connectivity. The resultant network should then be able to support auditory speech repetition. We plan to pursue this hypothesis in further work.

## Material and methods

### Participants

A total of 59 participants were included in this study. Participant details are provided in Table 2A. All participants were native English speakers, right-handed (assessed with the Edinburgh handedness inventory ^41^) neurologically intact and reported normal or corrected-to-normal vision and hearing. The study was approved by the London Queen Square Research Ethics Committee. All participants gave written informed consent before participation and were compensated £10 per hour for their time.

### Experimental paradigm

The current study focused on brain activation elicited when our participants were repeating heard words or pseudowords in different scanning runs. The words included object names that had an average of 1.68 syllables (range = 1 to 4) and an average duration of 0.65s (standard deviation = 0.08s). For pseudowords, the average syllables were 1.50 (range = 1 to 4) and an average duration of 1.45s (standard deviation = 0.15s). In each scanning run of 3.4 minutes, 40 words or pseudowords were presented sequentially with 4 blocks of 10 stimuli (25 seconds per block) interspersed with 16 seconds of rest (see Table 2B for further details).

In addition to word and pseudoword repetition, all participants performed 11 other conditions that are not part of the current study (see Paradigm 2 in ^42^). Crucially, the order of all conditions, the content of the stimuli and the presentation parameters were identical for all participants, therefore inter-participant variability in brain activation cannot be explained by any of these factors.

### Data acquisition and analysis

Functional MRI (fMRI) data were acquired on a 3T Trio scanner (Siemens Medical Systems) using a 12-channel head coil and a gradient-echo EPI sequence with 3 × 3 mm in-plane resolution (repetition time/echo time/flip angle: 3080 ms/30 ms/90°, extended field of view = 192 mm, matrix size = 64 × 64, 44 slices, slice thickness = 2 mm, and interslice gap = 1 mm). Structural MRI data were high-resolution T1-weighted images, acquired on the same 3T scanner using a 3D modified driven equilibrium Fourier transform sequence ^43^: TR/TE/TI = 7.92 ms/2.48 ms/910 ms, Flip angle = 16, 176 slices, voxel size = 1×1×1 mm3.

All data processing and analyses were performed with the Statistical Parametric Mapping (SPM12) software package (Wellcome Centre for Human Neuroimaging, London UK; http://www.fil.ion.ucl.ac.uk/spm/). All functional volumes were spatially realigned, unwarped, normalised to MNI space using a standard normalisation-segmentation procedure, and smoothed with a 6 mm full-width half-maximum isotropic Gaussian kernel, with a resulting voxel size of 3 × 3 × 3 mm.

### Brain region selection

The region of interest (ROI) selection process involved two steps. First, we defined the anatomical boundaries of each of our four regions of interest using the Brainnetome atlas ^44^ (Figure 1a). We selected regions Te1.0 and Te1.2 for the primary auditory cortex (A1), the rostroposterior STS subregion for Wernicke’s area (pSTS), the dorsal pOp subregion for Broca’s Area (dpOp), and the face (including the mouth) subregion for the primary motor cortex (M1-f). These choices were guided by the fMRI findings from an independent group of 25 neurologically intact participants who performed the same word and pseudoword repetition tasks as reported in Hope et al. (2014). Additionally, to evaluate functional degeneracy, we investigated the effects of exchanging the dorsal pOp subregion with the ventral pOp subregion (vpOp) and the face motor control subregion with the tongue and larynx motor control subregion (M1-tl) from the Brainnetome atlas. This resulted in 4 different (sub)regional configurations per subject for both word and pseudoword repetition (8 configurations per subject in total). The region borders were determined using a probability threshold of 50%: i.e., the anatomical localisation of the regions was consistent for at least 50% of the neurologically intact participants who contributed to the atlas construction. These probability thresholds are within the range used in previous studies ^29, 45, 46, 47^.

Second, we searched for the peak response during word and pseudoword repetition within each anatomically defined ROI (Figure 1a; Table 3) in each of the 59 participants. Separate time series of activation during the word and pseudoword repetition tasks were extracted from the peak coordinates for each participant. This ensured that effective connectivity between regions was estimated where activation was most robust for each participant, within a given ROI. In other words, we used each subject’s functional anatomy to define ROI specific responses. In each region, group activation during both word and pseudoword repetition was significant at voxelwise p<0.05 family-wise-error-corrected (using random field theory) for multiple comparisons across the whole brain (t-scores are reported in Table 3). Comparison of the coordinates for the peak response in each participant individually relative to that for the full group revealed that the mean distance was 0.74 mm (range = 0.37-1.56; standard deviation = 0.83).

**Table 3.**
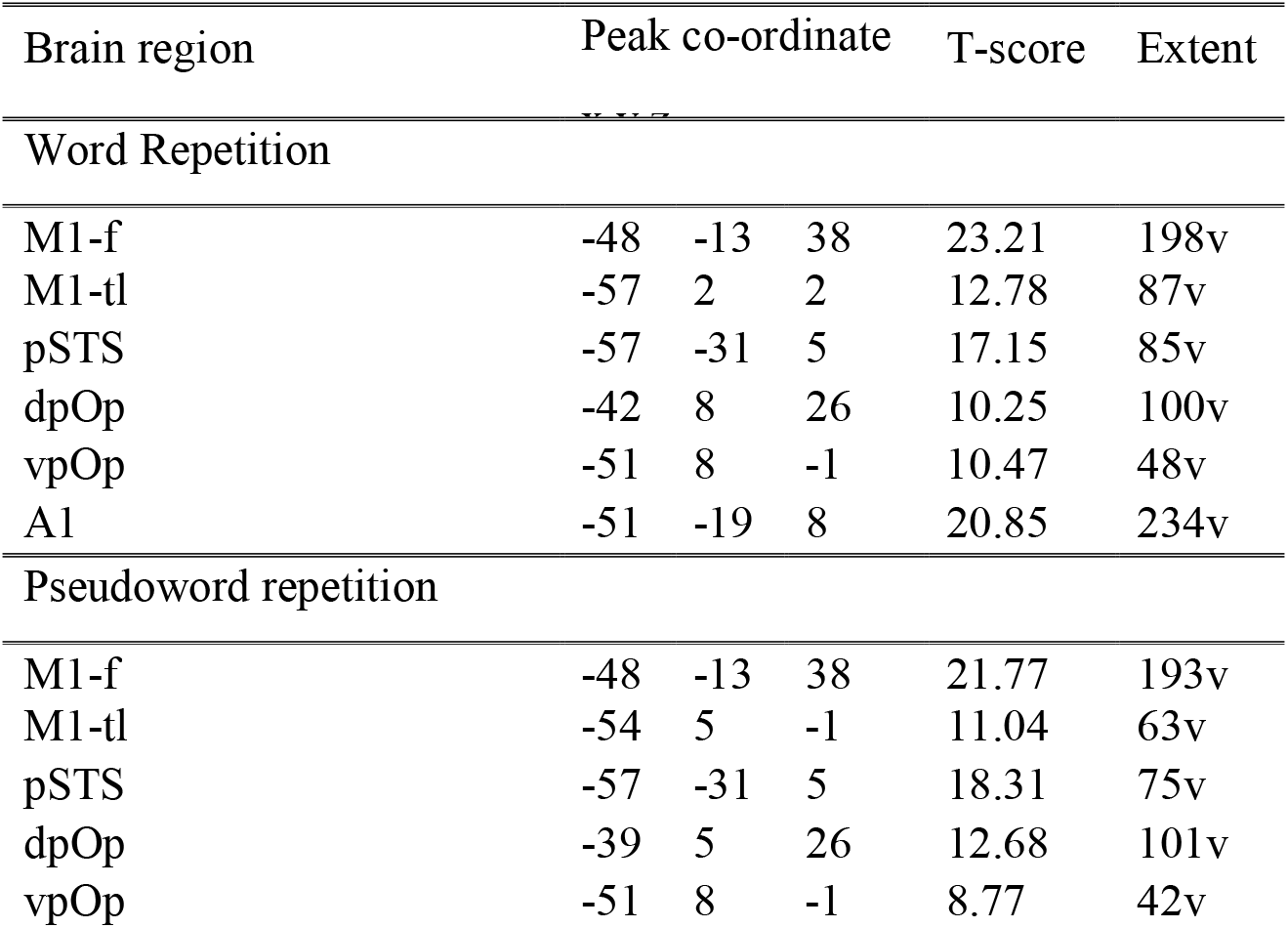
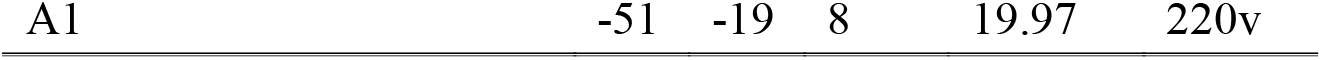
Effects reported have been thresholded at voxelwise p < 0.05 FWE-corrected.

### Dynamic causal modelling

Effective connectivity among four ROIs was estimated using dynamic causal modelling (DCM) ^14, 48^ as implemented in SPM12. DCM is a hypothesis-driven framework for investigating models of effective connectivity in a network of interconnected neuronal populations using approximate Bayesian inference. It characterises the brain as a nonlinear dynamical system of interconnected neuronal populations whose directed connection strengths may be modulated by endogenous activity or external perturbations. Briefly, the model consists of a neuronal model, and a forward model, that describes how activity at the neuronal level translates into observed signals (Figure 5a). DCM strives for a mechanistic explanation of experimental measures of brain activity in terms of directed intrinsic (self) and extrinsic (between region) connectivity. See 11 for a detailed overview.

**Figure 5.**
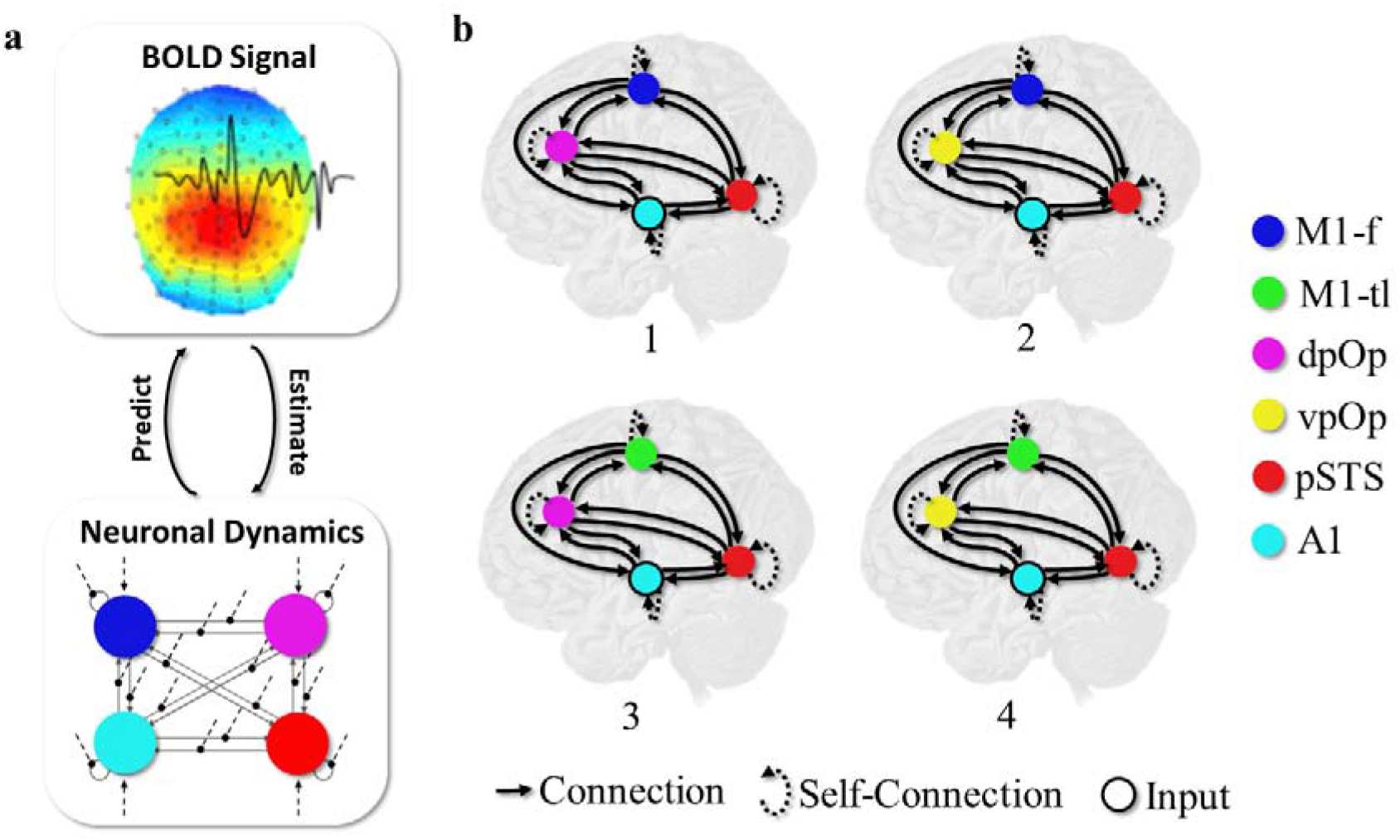
(a) Graphical illustration of DCM, and (b) Schematic view of the subject-level DCM. (a) presents a graphic illustration of the DCM for generative model: BOLD (blood oxygenation level-dependent) signal represents the observed fMRI data, Neuronal Dynamics represent the neural state dynamics and arrows between denote the forward model (i.e., predict) and inverse model (i.e., estimate). (b) presents the four different DCMs estimated. Each model comprised 4 regions with 15 connections, including the 4 self-connections. However, different subregional configurations lead to 4 different models (1-4): 1 (M1-f and dpOp), 2 (M1-f and vpOp), 3 (M1-tl and dpOp) and 4 (M1-tl and dpOp).

In the current study, we used two model parameters: (i) input parameters that identify which region was responding to external stimuli, here the primary auditory cortex; and (ii) the effective connectivity changes that occur among regions, as participants alternate between repetition and rest. These parameters were estimated at the neuronal level and the coupling between regions does not necessarily reflect the existence of direct (e.g., monosynaptic) connections.

For the technically savvy reader, we deliberately kept the analysis, and our interpretations, simple by separately estimating the average effective connectivity during each repetition task. In other words, our DCM models do not estimate the modulation of effective connectivity due to different experimental condition i.e., word or pseudoword. Therefore, all interpretations of reported estimates should be read as average effectivity connectivity, and not the rate of change in the effective connectivity due to modulatory inputs.

### Participant-level DCM

We now turn to the model specification. For each participant (and subregional configuration), we specified the model as defined in (Figure 5b): (i) the driving input was from the primary auditory cortex (A1), (ii) A1 was connected to all regions except the primary motor cortex M1 (i.e., M1-f or M1-tl) given anatomical constraints, (iii) pSTS was connected to pOp (i.e., dpOp or vpOp), and (iv) M1 (i.e., M1-f or M1-tl) was defined as the output region that received inputs from either pSTS, pOp or both. Briefly, all specified connections were both forward and backward, and they could be either excitatory or inhibitory. Importantly, our specification formulated pSTS and pOp at a similar level in the structural hierarchy because they were both connected to the input in A1 and the output in M1. Moreover, this allowed us to estimate whether pOp was higher (or lower) than pSTS within the functional hierarchy

We also specified an inhibitory self-connection for each region (which enables us to measure a region’s sensitivity to its inputs). Changes in these self-connections can be regarded as a reflection of excitatory-inhibitory balance within each region ^22^. The parameters are set to be negative (default is -0.5Hz) to preclude run-away excitation in the network ^11^. Accordingly, positive self-connection estimates are indicative of inhibition and negative self-connections are indicative of excitation.

All model parameters and their posterior probabilities were estimated, with Bayesian inversion, using variational Laplace ^14^, an automatic variational procedure under Gaussian assumptions about the form of the posterior. The participant-level specification was separately estimated for the different subregional configurations per participant for both word and pseudoword repetition, i.e., 8 DCM model estimations per participant in total.

### Group-level DCM

We evaluated group effects and between-participant variability on parameters using the Parametric Empirical Bayes (PEB) model ^12^. The resultant hierarchical model quantifies the estimated connection strengths, and their uncertainty, from the participant to the group level. Having estimated the group-level parameters (e.g., group-average effective connection strengths), we used Bayesian model comparison to test hypotheses for alternative models of effective connectivity during word and pseudoword repetition separately. The alternative models were generated by switching parameters on and off using an automatic grid search ^12^. Explicitly, the group-level DCM estimates were evaluated across the 8 subregional configurations (4 for word repetition and 4 for pseudoword repetition). For each configuration, we evaluated the model evidence across 256 separate models where each model represented the removal of a particular parameter or effective connection e.g., from A1 to pSTS or pSTS to M1. Finally, the model with the highest model evidence was selected for each of the 8 subregional configurations.

## Supplementary Material

**Supplementary Figure 1.**
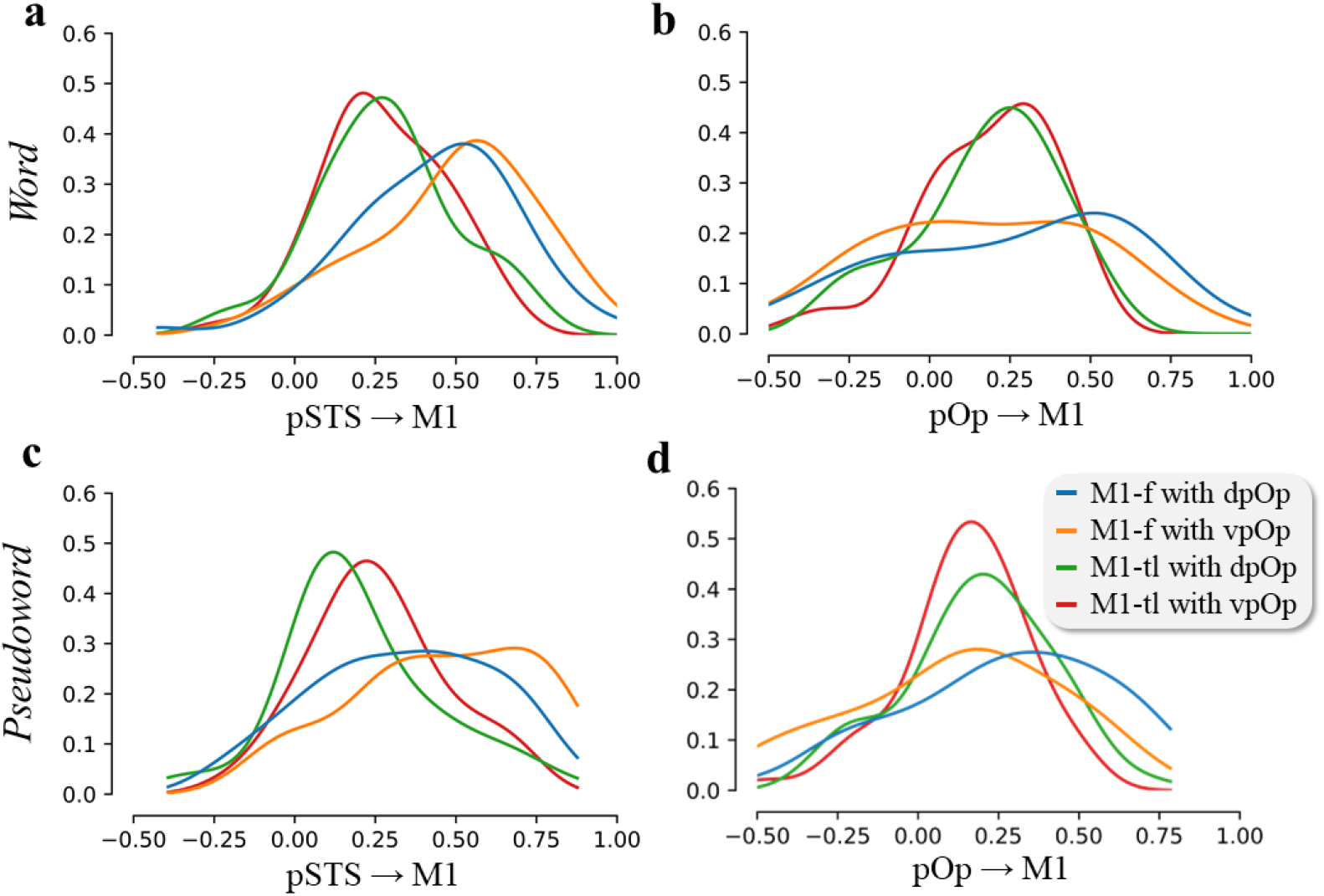
Individual-level effective connectivity from pOp and pSTS to M1. The two panels in the first row (a) summarise connections across the different subregional configurations for word repetition, and the two panels in the second row (b) summarise connections across the different subregional configurations for pseudoword repetition: blue (M1-f and dpOp), orange (M1-f and vpOp), green (M1-tl and dpOp) and red (M1-tl and dpOp). The first panel in each row presents the sample density for individual connections from pSTS to M1 and the second panel in each row presents the sample density over the connections from pOp to M1.

**Supplementary Figure 2.**
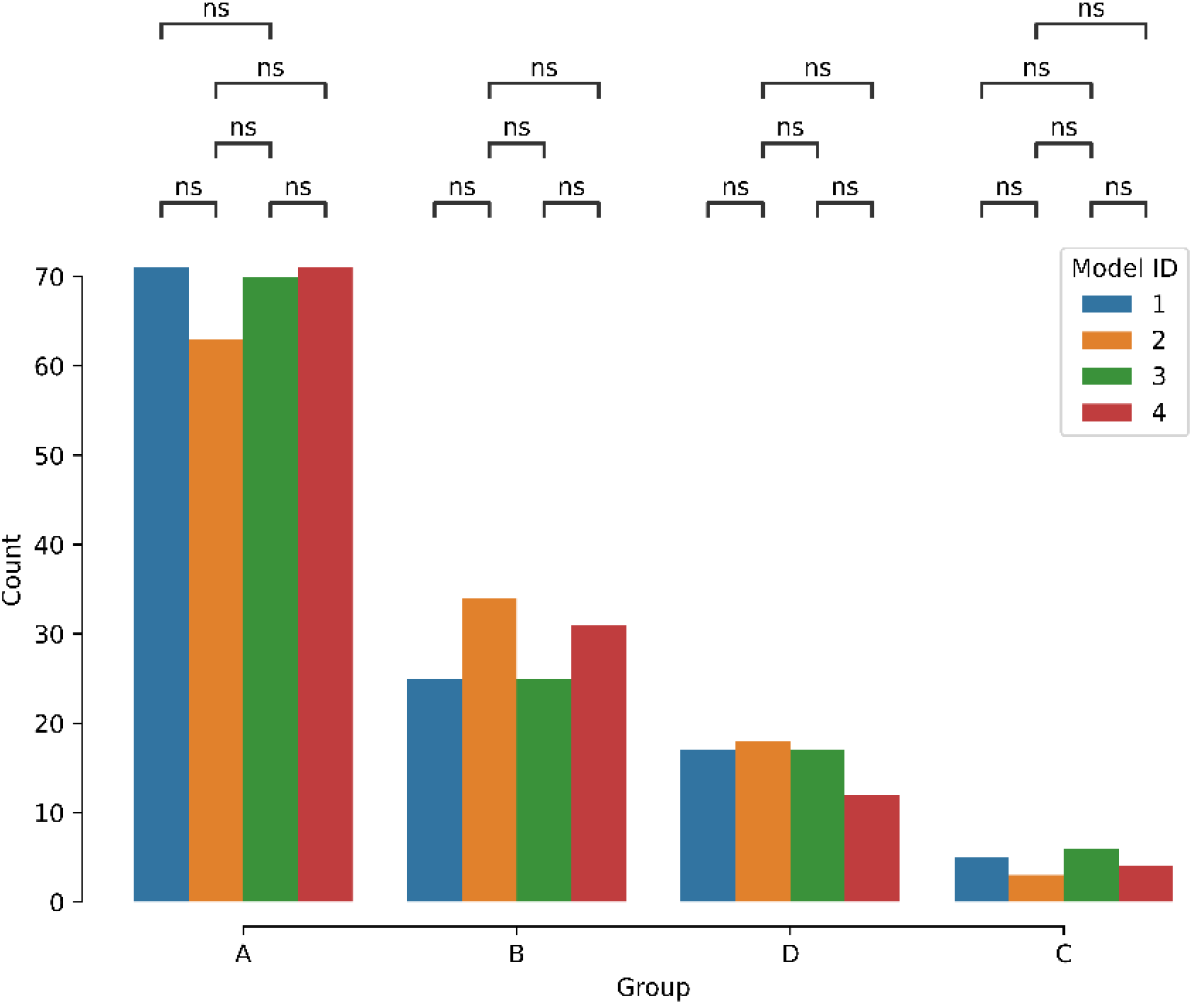
Variability in group membership. The bar plot reports the differences in group membership across the different model configurations: 1 (M1-f and dpOp), 2 (M1-f and vpOp), 3 (M1-tl and dpOp) and 4 (M1-tl and dpOp). Here, the x-axis is the group, and the y-axis reports the total number of models in that group. We found no significant differences (with p-value of 1.00) between word and pseudoword repetition using a two-sided Mann-Whitney-Wilcoxon test with Bonferroni correction.

**Supplementary Figure 3.**
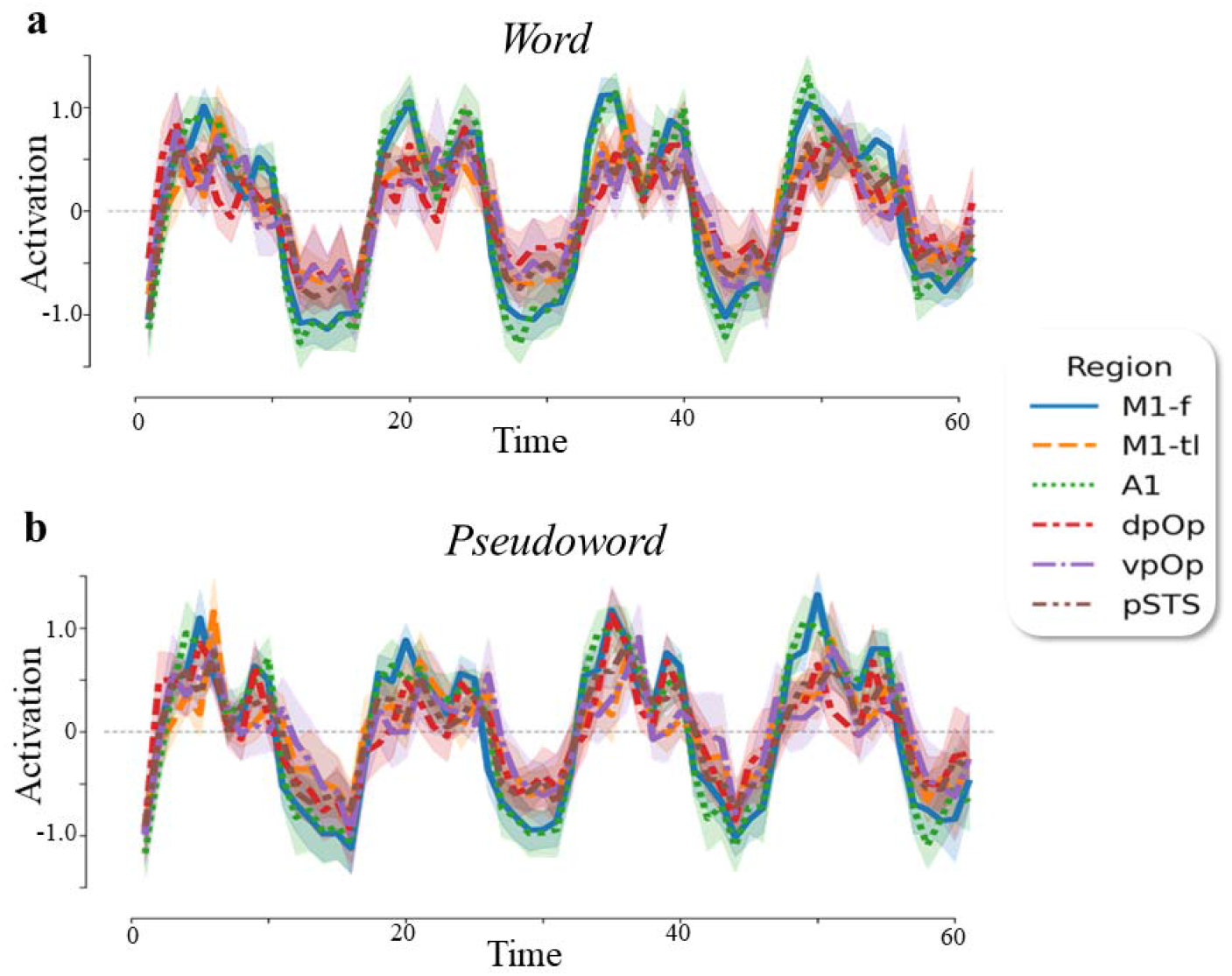
Activation in each subregion of interest (ROI) during word and pseudoword repetition. The plots indicate activation in each ROI over the trial duration during word (a) and pseudoword repetition (b). Specifically, x-axis is the first principal component of the pre-whitened, high-pass filtered and confounded corrected timeseries for each ROI. The y-axis plots brain activation in each region: M1-f (blue), M1-tl (orange), dpOp (red) vpOp (purple), pSTS (brown) and A1 (green).

**Supplementary Table 1.** Group membership across the 59 subjects for each M1 (M1-f or M1-tl) and pOp (dpOp or vpOp) region during word and pseudoword repetition. Here, A had positive connections from pOp to M1 and pSTS to M1; B had positive connections from pSTS to M1 but not from pOp to M1; C had positive connections from pOp to M1 but not from pSTS to M1, and D denotes the posterior probability of < 0.75 i.e., no significant connections from both pOp and pSTS to M1. Moreover, consistency with the neurological model requires excitatory connectivity from pSTS to pOp, and pOp to M1 but not pSTS to M1. Only estimated connections with a high posterior probability (i.e., > 0.75) were included. * denotes evidence for the neurological model. Here, 1-4 denotes the different subregional configurations: 1 (M1-f and dpOp), 2 (M1-f and vpOp), 3 (M1-tl and dpOp) and 4 (M1-tl and dpOp).

|  | Word |  |  |  | Pseudowords |  |  |  |
| --- | --- | --- | --- | --- | --- | --- | --- | --- |
| ID | 1 | 2 | 3 | 4 | 1 | 2 | 3 | 4 |
| C010 | A | A | A | A | A | A | A | A |
| C012 | A | A | A | A | A | B | D | B |
| C029 | A | A | A | A | A | B | A | A |
| C048 | B | B | B | B | D | B | B | B |
| C050 | B | A | A | A | A | A | A | A |
| C052 | A | A | D | A | A | A | A | B |
| C065 | A | A | A | D | B | B | B | B |
| C067 | A | D | D | D | A | A | C | D |
| C068 | A | A | A | A | A | A | A | A |
| C069 | B | B | A | A | B | B | D | D |
| C071 | D | A | C | A | A | A | A | A |
| C077 | A | D | A | A | A | A | A | A |
| C078 | A | B | A | A | A | A | A | A |
| C110 | A | A | A | A | A | A | A | A |
| C001 | D | B | D | D | D | A | D | D |
| C006 | B | A | B | A | A | A | A | A |
| C008 | D | A | B | A | A | D | A | A |
| C042 | A | A | A | A | A | A | A | A |
| C043 | A | A | A | A | A | A | A | A |
| C049 | A | A | A | D | B | B | A | A |
| C055 | A | D | B | D | B | B | B | B |
| C063 | A | A | B | D | A | A | A | A |
| C064 | B | B | A | A | B | B | D | A |
| C066 | B | B | D | C* | A | D | A | A |
| C079 | A | A | D | B | D | D | D | C |
| C081 | A | A | A | A | D | D | D | D |
| C087 | D | A | B | A | A | D | A | A |
| C160 | C* | A | A | A | D | D | D | D |
| C161 | B | B | D | D | A | A | A | D |
| C059 | D | A | D | A | A | B | A | A |
| C070 | D | A | A | A | A | A | A | A |
| C073 | B | C | B | B | D | A | C* | C |
| C074 | B | B | D | D | A | D | A | A |
| C075 | A | B | A | A | D | B | A | A |
| C080 | A | A | A | A | A | A | D | A |
| C083 | B | B | A | B | B | B | B | B |
| C111 | A | D | A | A | A | D | A | A |
| C112 | B | B | D | D | A | A | D | D |
| C167 | A | A | B | D | B | C | D | A |
| C168 | A | A | A | A | C* | A | B | A |
| C169 | D | C* | A | D | B | D | A | A |
| C170 | B | B | A | A | A | A | B | A |
| C041 | A | A | A | A | A | A | A | A |
| C044 | B | B | A | A | B | B | A | D |
| C045 | C* | B | A | D | A | A | A | A |
| C046 | A | D | A | A | A | A | D | A |
| C047 | A | A | A | D | A | A | D | D |
| C053 | A | B | A | A | C* | A | A | B |
| C056 | A | A | A | A | C* | D | D | D |
| C057 | B | D | A | A | D | B | A | A |
| C058 | A | A | A | A | A | A | A | A |
| C060 | A | A | C | A | B | B | B | A |
| C061 | A | D | C* | D | A | A | A | A |
| C062 | A | B | A | D | A | B | D | B |
| C108 | A | A | A | A | A | A | C* | C |
| C104 | A | A | A | A | A | B | A | D |
| C040 | A | A | A | D | D | D | D | A |
| C051 | B | B | B | D | A | A | D | D |
| C054 | A | B | A | D | D | A | B | D |

**Supplementary Table 2A.**
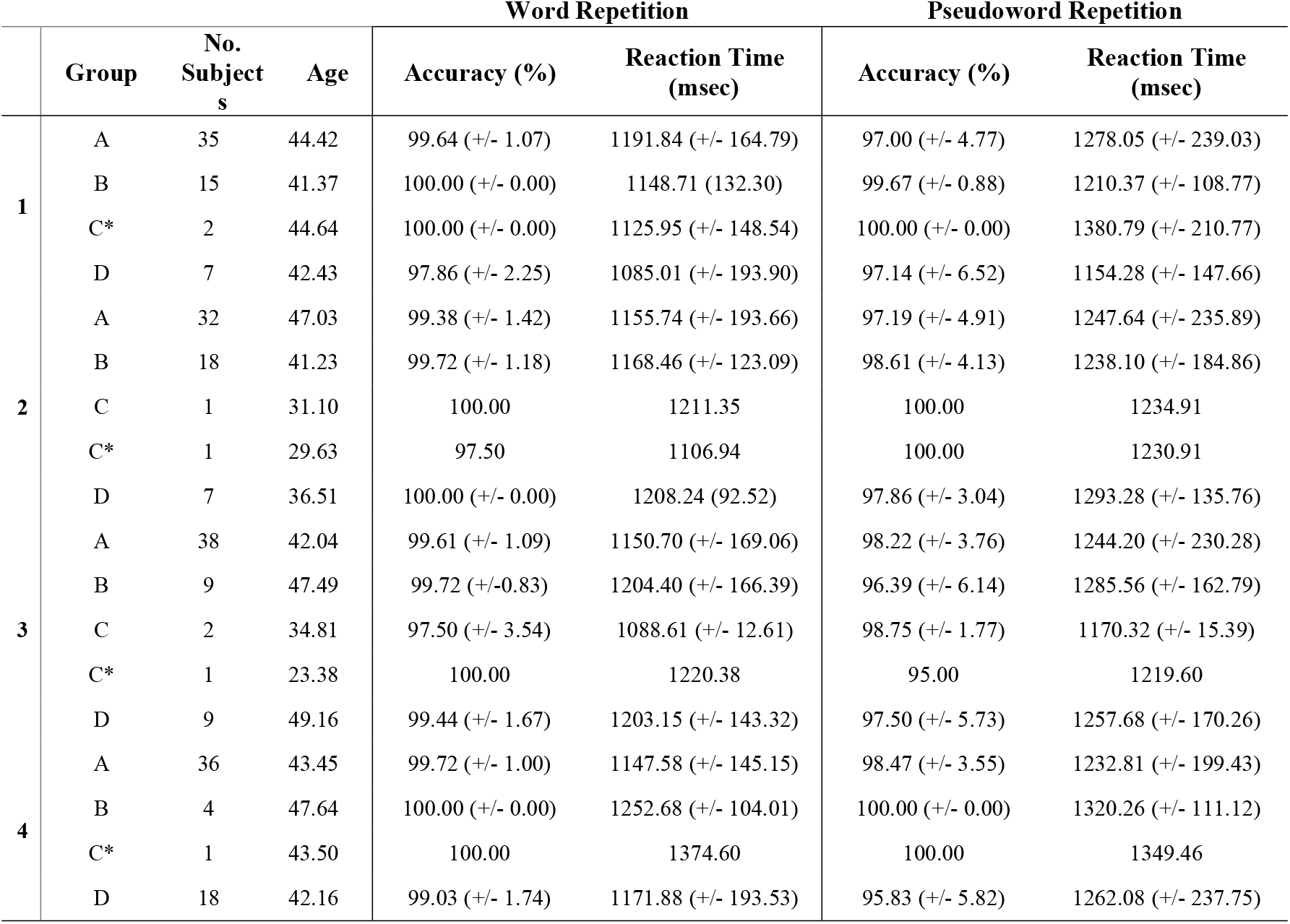
Summary of participant data by group, for each word repetition DCM subregional configuration. Here, 1-4 denotes the different subregional configurations: 1 (M1-f and dpOp), 2 (M1-f and vpOp), 3 (M1-tl and dpOp) and 4 (M1-tl and dpOp). Here, A had positive connections from pOp to M1 and pSTS to M1; B had positive connections from pSTS to M1 but not from pOp to M1; C had positive connections from pOp to M1 but not from pSTS to M1, and D denotes the posterior probability of < 0.75 i.e., no significant connections from both pOp and pSTS to M1. Additionally, C* denotes consistency with the neurological model.

**Supplementary Table 2B.**
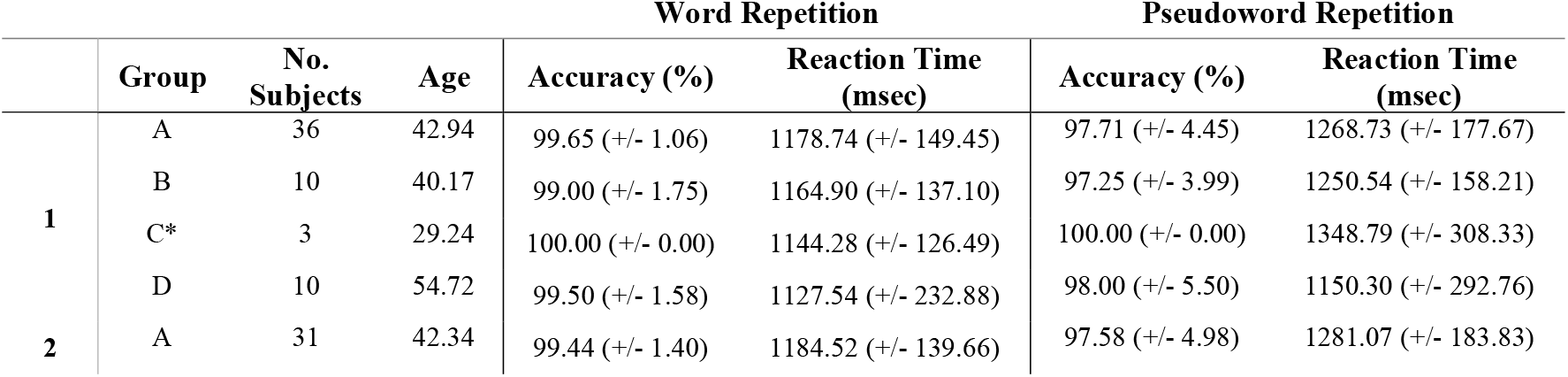
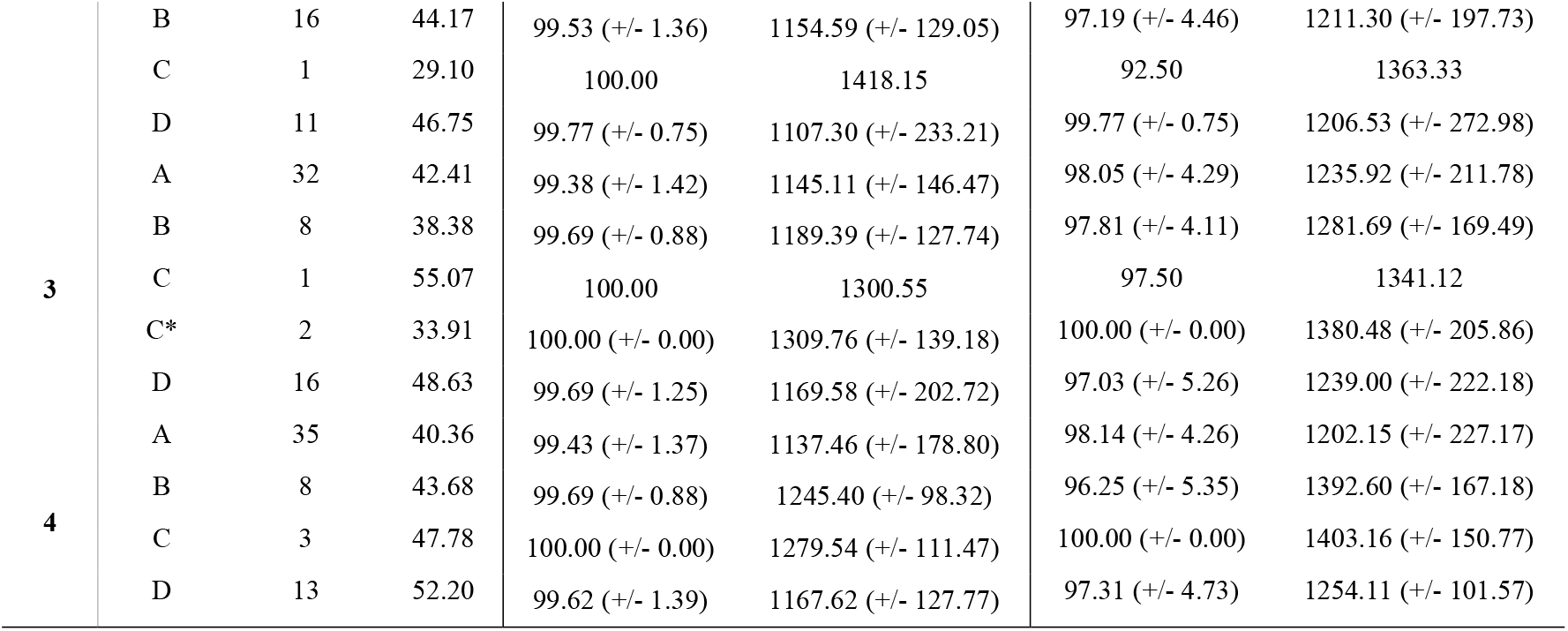
Summary of participant data by group, for each pseudoword repetition DCM subregional configuration. Here, 1-4 denotes the different subregional configurations: 1 (M1-f and dpOp), 2 (M1-f and vpOp), 3 (M1-tl and dpOp) and 4 (M1-tl and dpOp). Here, A had positive connections from pOp to M1 and pSTS to M1; B had positive connections from pSTS to M1 but not from pOp to M1; C had positive connections from pOp to M1 but not from pSTS to M1, and D denotes the posterior probability of < 0.75 i.e., no significant connections from both pOp and pSTS to M1. Additionally, C* denotes consistency with the neurological model.

